# Evolution of density-dependent movement during replicated experimental range expansions

**DOI:** 10.1101/114330

**Authors:** Emanuel A. Fronhofer, Sereina Gut, Florian Altermatt

**Author notes:** Correspondence Details Emanuel A. Fronhofer Eawag: Swiss Federal Institute of Aquatic Science and Technology Department of Aquatic Ecology Überlandstrasse 133 CH-8600 Dübendorf.

## Abstract

Range expansions and biological invasions are prime examples of non-equilibrium systems that are likely impacted by rapid evolutionary changes. As a spatial process, range expansions are driven by dispersal and movement behaviour. While it is widely accepted that dispersal and movement may be context-dependent, for instance density-dependent, and best represented by reaction norms, the evolution of density-dependent movement during range expansions has received little experimental attention. We therefore tested current theory predicting the evolution of increased movement at low densities at range margins using highly replicated and controlled range expansion experiments across multiple genotypes of the protist model system *Tetrahymena thermophila*. Although rare, we found evolutionary changes during range expansions even in the absence of initial standing genetic variation. Range expansions led to the evolution of negatively density-dependent movement at range margins. In addition, we report the evolution of increased competitive ability and concurrently decreased population growth rates in range cores. Our findings highlight the importance of understanding movement and dispersal as evolving reaction norms and plastic life-history traits of central relevance for range expansions, biological invasions and the dynamics of spatially structured systems in general.

**Data accessibility:** The data will be made available at Dryad (http://dx.doi.org/xxxxx). Video analysis scripts can be found on GitHub (https://github.com/efronhofer/analysis_script_bemovi).

## Introduction

Species’ ranges and the distribution of taxa on earth have long been studied only from an ecological point of view and put in the context of local climatic and environmental conditions. Recent theoretical work and empirical case examples however, have expanded upon this work by including an evolutionary perspective. Such an approach is needed because rapid evolutionary changes are thought to be especially relevant under transient and non-equilibrium conditions inherent to species range dynamics and population spread (Excoffier *et al.*, 2009; Hanski, 2012; Kubisch *et al.*, 2014).

The importance of an evolutionary and eco-evolutionary perspective for understanding the dynamics of species ranges becomes evident when considering the main drivers of range dynamics: dispersal and reproduction. These life-history traits not only have a genetic basis (Bonte & Dahirel, 2017; Saastamoinen *et al.*, submitted) and can be subject to evolutionary change, but ample theoretical evidence suggests that the evolution of dispersal and reproduction may also have important effects on the dynamics of species ranges (e.g. reviewed in Kubisch *et al.*, 2014). For instance, dispersal during range expansions is predicted to evolve to higher rates at range margins, a phenomenon called ‘spatial selection’ (Phillips *et al.*, 2010). Spatial selection relies on a simple ecological filter effect: if there is variation in dispersal ability, fast individuals will automatically be spreading more towards the range margin and colonize previously empty patches (see also Haag *et al.*, 2005). As a consequence, populations at the range margin have a higher proportion of more dispersive individuals that will mate with each other. In combination with fitness advantages due to low intraspecific competition at the range margin, this spatial assortment can lead to the evolution of increased dispersal and movement abilities. These evolutionary changes have important consequences for (macro)ecological patterns: range expansions and biological invasions are predicted to proceed faster if dispersal evolves (Perkins *et al.*, 2013). Theoretical predictions regarding the evolution of reproduction have also been made, especially in the context of life-history trade-offs between dispersal, reproduction and competitive ability (Burton *et al.*, 2010; Fronhofer & Altermatt, 2015; Fronhofer *et al.*, 2017). Since dispersal is costly (Bonte *et al.*, 2012), models generally assume that either reproduction or competitive ability have to decrease if dispersal increases.

While there is accumulating comparative empirical evidence linked to spatial selection (e.g. Thomas *et al.*, 2001; Phillips *et al.*, 2006; Lombaert *et al.*, 2014), experimental confirmation is more scare. This scarcity is mainly due to the obvious challenges associated with studying how evolution impacts macroecological patterns experimentally. However, a recently increasing use of (small) model organisms has been spurring the study of experimental range expansions and population spread dynamics. As a consequence, range expansions are currently under intense scrutiny from an experimental ecological (e.g. Melbourne & Hastings, 2009; Giometto *et al.*, 2014, in press) and evolutionary point of view (e.g. Hallatschek *et al.*, 2007; Fronhofer & Altermatt, 2015; Gralka *et al.*, 2016; Williams *et al.*, 2016; Wagner *et al.*, 2017; Ochocki & Miller, 2017; Weiss-Lehman *et al.*, 2017; Fronhofer *et al.*, 2017). A result of these concerted efforts is the experimental demonstration that range expansions indeed may select for increased dispersal rates and distances at the rage margin (in protists: Fronhofer & Altermatt 2015; Fronhofer *et al.* 2017; in insects: Ochocki & Miller 2017; Weiss-Lehman *et al.* 2017 and in plants: Williams *et al.* 2016), which can lead to increased range expansion speeds.

We build on these studies and address in more detail two aspects that have received less attention so far: Firstly, we explore evolutionary change during range expansions of populations initially harbouring no standing genetic variation. Understanding the consequences of depleted genetic variation may be especially relevant in the context of biological invasions, since genetic variation may be lost due to bottlenecks associated to the invasion process itself, which potentially constrains the success of invasions and evolutionary dynamics (for a critical discussion, see Roman & Darling, 2007; Dlugosch & Parker, 2008). The literature on biological invasions also suggests that successful range expansions should be overall rare, regardless of whether evolutionary changes occur or not. For example, the classic, empirically derived ‘tens rule’ discussed by Williamson & Fitter (1996) states that one tenth of imported species become introduced, one tenth of these species will become established and again only one tenth will successfully spread.

Secondly, we highlight that movement is context-dependent (Clobert *et al.*, 2009; Fronhofer *et al.*, 2017; Weiss-Lehman *et al.*, 2017), that is, a reaction norm. The most well-known dispersal reaction norm captures dispersal as a function of conspecific density (e.g. Pennekamp *et al.*, 2014; Fronhofer *et al.*, 2015b), as the spatio-temporal distribution of densities is an important proximate and ultimate factor determining dispersal strategies (Bowler & Benton, 2005; Ronce, 2007). Current theory predicts that range expansions do not only select on the mean dispersal trait, but rather on the shape of the reaction norm (Travis *et al.*, 2009). Dispersal is predicted to increase even at low densities which makes the reaction norm less positively density-dependent at the range margin (Travis *et al.*, 2009). While density-dependent dispersal will optimize fitness in (quasi)equilibrium environments, such as the range core (Bowler & Benton, 2005), local conspecific density does not provide relevant information at the range margin: spatial selection makes dispersal highly advantageous even if local densities at the range margin are low. The evolution of increased dispersal at low densities at the range margin may have important consequences for range expansion dynamics: theory shows that density-dependent dispersal may slow down range expansions (Best *et al.*, 2007) but can ultimately lead to wider ranges (Kubisch *et al.*, 2011), to name but two examples.

Here, we experimentally tested whether and how range expansions impact the evolution of density-dependent movement (as a proxy of dispersal) and potentially change the shape of the density-dependent movement reaction norm. We predicted movement to evolve towards higher values at low densities making the reaction less positively or even negatively density-dependent at the range margin (Travis *et al.*, 2009; Weiss-Lehman *et al.*, 2017). We tested this prediction using experimental evolution and a total of 48 replicate range expansions of 14 clonally reproducing *Tetrahymena thermophila* strains, as well as the mix of these strains. Due to the lack of initial standing genetic variation in the replicates with single clonal strains and the low population densities characteristic of expanding range margins, we predicted consistent evolutionary differentiation between range core and margin populations to be rare and emerge only in a minority of our replicates. In addition, successful range expansions, whether including evolutionary change or not, can be expected to be rare overall, as suggested by comparative empirical evidence of biological invasions (Williamson & Fitter, 1996). We recorded movement behaviour and changes in life-history traits, namely population growth rate and competitive ability, to investigate potential concurrent evolutionary changes (Bonte & Dahirel, 2017).

## Material and Methods

### Model organism

We used the freshwater ciliate *Tetrahymena thermophila* (Cassidy-Hanley, 2012) as our model organism. It is, together with its sister species (*Tetrahymena pyriformis*), commonly and successfully employed in both ecological (e.g., Giometto *et al.*, 2014; Fronhofer *et al.*, 2015b) and evolutionary contexts (e.g., Fjerdingstad *et al.*, 2007; Friman *et al.*, 2011; Pennekamp *et al.*, 2014; Hiltunen & Becks, 2014; Fronhofer & Altermatt, 2015) and is known to show density-dependent movement and dispersal (Pennekamp *et al.*, 2014; Fronhofer *et al.*, 2015b). *Tetrahymena thermophila* can reproduce both sexually (if a cell from another mating type is present and mating is induced, e.g. by starvation, which was not the case in our experiments) and clonally (Cassidy-Hanley, 2012). We used a total of 14 clonally reproducing strains that were isolated as single cells shortly prior to the experiment from 7 mating types originally obtained from Cornell University’s Tetrahymena Stock Center (Stock ID: MTP01). The 14 clonally reproducing strains used here differ in attributes ranging from population growth rates and equilibrium densities (Tab. S1) to morphology and movement characteristics (Tab. S2).

*Tetrahymena thermophila* was kept in filtered (cellulose paper filters, 4–7 *μm*; Whatman) and autoclaved protist pellet medium (0.46 *gL*^−1^; Protozoan pellets, Carolina Biological Supply) under constant temperature (20*°C*) and light conditions (Osram Biolux 30W). Initially, we supplemented the medium with a mix of three bacterial species (*Serratia fonticola*, *Bacillus subtilis* and *Brevibacillus brevis* ; obtained from Carolina Biological Supply) at approx. 5% of their equilibrium density as a food source. During the experiment we replaced 2 mL out of 15 mL medium in each microcosm with bacterial cultures at equilibrium density (approx. 1 week old) in order to prevent adaptation of the bacteria and co-evolutionary dynamics from occurring. For further methodological details see Altermatt *et al.* (2015).

### Experimental setup and microcosm landscapes

We set up microcosm landscapes to study range expansions into initially uninhabited habitat by propagating range core and range margin populations as described by Fronhofer & Altermatt (2015). The microcosms landscapes consisted of two vials (20 mL conical vials, Sarstedt) connect by silicone tubing (inside diameter: 4 mm, VWR) and a stopcock (Discofix, B.Braun) to control for dispersal (total length: 6 cm). For a graphical overview of the experimental procedure see Fig. S1.

The first patch (start) was initially filled with the respective *T. thermophila* strain at equilibrium density and dispersal into the empty target patch (filled with 15 mL protist pellet medium with bacteria) was allowed for 4 h. Thereafter, the populations in the start and target patches were propagated to start patches in respectively new 2-patch systems. The population from the original start patch was subsequently taken as the range core. Dispersal was allowed for 4 h and only the individuals remaining in the start patch were further propagated. By contrast, the population from the original target patch represented the range margin. Dispersal was also allowed for 4 h. Thereafter, the population in the target patch was propagated to a new 2-patch system. This procedure, effectively tracking range core and expanding range margin populations, was repeated 3 times per week for 1 month.

During transfer to new 2-patch systems only 13 of the 15 mL of the range core were transferred and the remaining 2 mL were filled with bacterial cultures at equilibrium (approx. 1 week old) to avoid evolutionary changes in the bacterial populations. This procedure was not necessary for the range margin, as the individuals of the range margin had always just dispersed into medium with bacteria from the stock population.

We chose this experimental approach that tracks only range core and range margin for convenience and feasibility. Firstly, we can easily control resource levels and avoid loss of control and contaminations due to aging medium in long landscapes. Secondly, we have shown that, although the experimental procedure obviously prevents feedbacks between core and margin populations, our results are not altered if we indeed use long interconnected linear landscapes (Fronhofer *et al.*, 2017).

We followed a total of 48 range expansions, that is, pairs of range cores and margins. We replicated the range expansion of each of the 14 strain three times and additionally included six replicates of a mix (in equal proportions) of all strains. The latter range expansions using the mix of strains thus differs from the range expansions of the single clones in that they were started with standing genetic variation.

### Recording of population densities and movement data

During the experiment all microcosms were sampled at every transfer, that is three times per week. Data on population densities as well as movement metrics (e.g., velocity of the cells) were collected by video recording and automated analysis.

Using a Leica M205C stereomicroscope at a 16-fold magnification and an Orca Flash 4 camera (Hamamatsu) we recorded 20 s videos (25 frames per second) of an effective volume of 34.4 *μL* (sample height: 0.5 mm). Videos were subsequently analysed using the R environment for statistical computing (version 3.2.3) and a customized version of the ‘BEMOVI’ package (available on GitHub: https://github.com/efronhofer/bemovi) originally developed by Pennekamp *et al.* (2015). The ‘BEMOVI’ package uses the image processing software ImageJ to locate moving particles in the videos and to calculate population densities as well as movement metrics. Data are automatically filtered to exclude artifacts such as debris that are outside the size or movement range of the study species (detailed settings available on GitHub: https://github.com/efronhofer/analysis script bemovi). As movement velocity has been shown to correlate well with dispersal rates in this system (Fronhofer & Altermatt, 2015; Fronhofer *et al.*, 2015a) we will focus on velocity as a proxy for dispersal.

### Common garden experiments

In order to separate evolutionary from potential plastic intergenerational and environmental effects, movement behaviour of individuals from range core and range margin populations was measured not only during the evolution experiment, but also after the end of the experiment and a three-day common garden phase (doubling time for the strains of *T. thermophila* used here: approx. 3–12 h; see Tab. S1). The common garden was initiated with 1.5 mL samples obtained from the respective core and margin populations from the last day of the evolution experiment that were added to 13.5 mL medium with bacteria from the stock population to homogenize environmental conditions.

In addition, we kept a subset of the individuals analysed in this first common garden for an extended period in a common environment (total of 38 days). This allowed us to explore the long-term stability of potential evolutionary changes. Resources were not refreshed in order to create strong competition similar to stable range core populations.

### Estimating population growth rates and competition coefficients

Both, at the beginning of the experiment and after the common garden we used population growth experiments to estimate population growth rates (*r*_0_) as well as competition coefficients (*α*_*ii*_). These experiments were always started with 1 mL of the respective strain (1 week old for the beginning; from the common garden for the end) and 14 mL medium with bacteria from the stock population. We then followed population densities and movement over the course of 9 days, measuring twice in the first two days and then once on days 2, 3, 4, 7 and 9.

Population growth rates (*r*_0_) and competition coefficients (*α*_*ii*_) were estimated by fitting the logistic growth model according to Verhulst (1838)

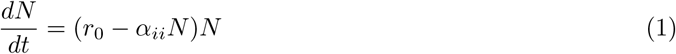

where *N* is the population size, to the data obtained from the growth experiments. The equilibrium density is obtained as 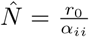. We used the R environment for statistical computing (version 3.3.2) to solve (function ‘ode’; ‘deSolve’ package) and subsequently fit the model by minimizing residuals using the Levenberg-Marquardt algorithm (function ‘nls.lm’; ‘minpack.lm’).

### Density-dependent movement

The data obtained from the growth experiments were also used to assess potential evolutionary changes in density-dependent movement (see also Fronhofer *et al.*, 2015b), that is, velocity as a function of relative population density 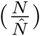. Correlating population densities and movement may be challenging: for instance, low densities due to declining or growing populations may lead to very different movement decisions, as protists likely base movement decisions on chemical cues (Fronhofer *et al.*, 2015b). We therefore determined whether there was an indication for a decay phase towards the end of the growth experiments by additionally fitting a logistic model that allows for decaying population densities. This growth model follows Eq. 1, however it assumes that *α*_*ii*_ = *α*_0_(1 + *β*(*t − t*_*crit*_)) if *t > t*_*crit*_ where *α*_0_ is a baseline competitive ability, *β* the rate at which competition increases over time (*t*) and *t*_*crit*_ the critical time point from which on the increase in competition (respectively decrease in equilibrium density) occurs. Based on these analyses, we excluded all movement data of day 7 of the growth experiment onwards as both mean and median values of *t*_*crit*_ estimates were approx. 7 days.

### Statistical analyses

All data were analysed statistically using the R environment for statistical computing (version 3.3.2) and linear mixed models (function ‘lme’; ‘nlme’ package). Response variables were appropriately transformed to satisfy model assumptions. Statistical analyses were performed across all strains simultaneously as the different genetic backgrounds represent the biologically relevant replicates in contrast to the technical replicates within each strain. This nested replication structure was accounted for in the random effects. The data are available at Dryad (http://dx.doi.org/xxxxx).

More specifically, movement velocity (log transformed) change during the experiment was analysed using individual based velocity data with ‘time’ within ‘replicate’ and ‘strain ID’ as a random effect structure to account for the non-independence of individuals measured in one sample as well as the temporal non-independence of the repeated measures through time. Following Crawley (2013), we started with the full model including an interaction between the two explanatory variables ‘time’ and ‘range position’ and subsequently simplified the model by removing explanatory variables to obtain the minimal adequate model. We used the ‘anova’ function and maximum likelihood (ML) estimates obtained with the ‘optim’ optimizer for model simplification. Subsequently, the minimal adequate model was refit using REML to obtain model estimates.

For the analysis of movement velocity (non-transformed) after the common garden phase the full model included ‘time’ (end of evolution experiment, common garden 1, common garden 2) and ‘range border position’ as interacting fixed effects as well as ‘time’ within ‘replicate’ within ‘strain ID’ as random effects. Model simplification was carried out as described above.

Density-dependent movement (log transformed) was analysed analogously using ‘population density’ and ‘population history’ (range border position and starting populations) as interacting fixed effects in the full model and ‘population density’ within ‘replicate’ within ‘strain ID’ as random effects. The analysis of population growth rates (*r*_0_; log transformed) and competition coefficients (*α*_*ii*_; inverse transformed) included only ‘population history’ (range border position and starting populations) as a fixed effect as well as ‘strain ID’ as a random effect.

Finally, potential differences between failed and successful range expansions, that is, experimental replicates in which the range margin failed to expand and went extinct versus those that kept up with our experimental treatment, were explored by correlating growth rates (*r*_0_; inverse transformed), competition coefficients (*α*_*ii*_; inverse transformed) as well as median and standard deviation of movement velocity of the starting populations (respectively log and inverse transformed) with invasion success (yes/ no). As a random effect we included ‘strain ID’.

## Results

### Range expansion success

As expected, successful range expansions were rare: out of 48 replicated range expansions only 4 replicates belonging to three clonal strains (Fig. 1; upper panel) successfully tracked our experimental treatment and expanded their range over the entire duration of the experiment. All other replicates of both clonal strains and the genetically mixed population only tracked the range expansion over part of the experimental duration and subsequently went extinct at the range margin. The four successfully expanding replicates did, at the start of the experiment, not statistically differ from all other stains based on their initial properties (Fig. S2).

**Figure 1:**
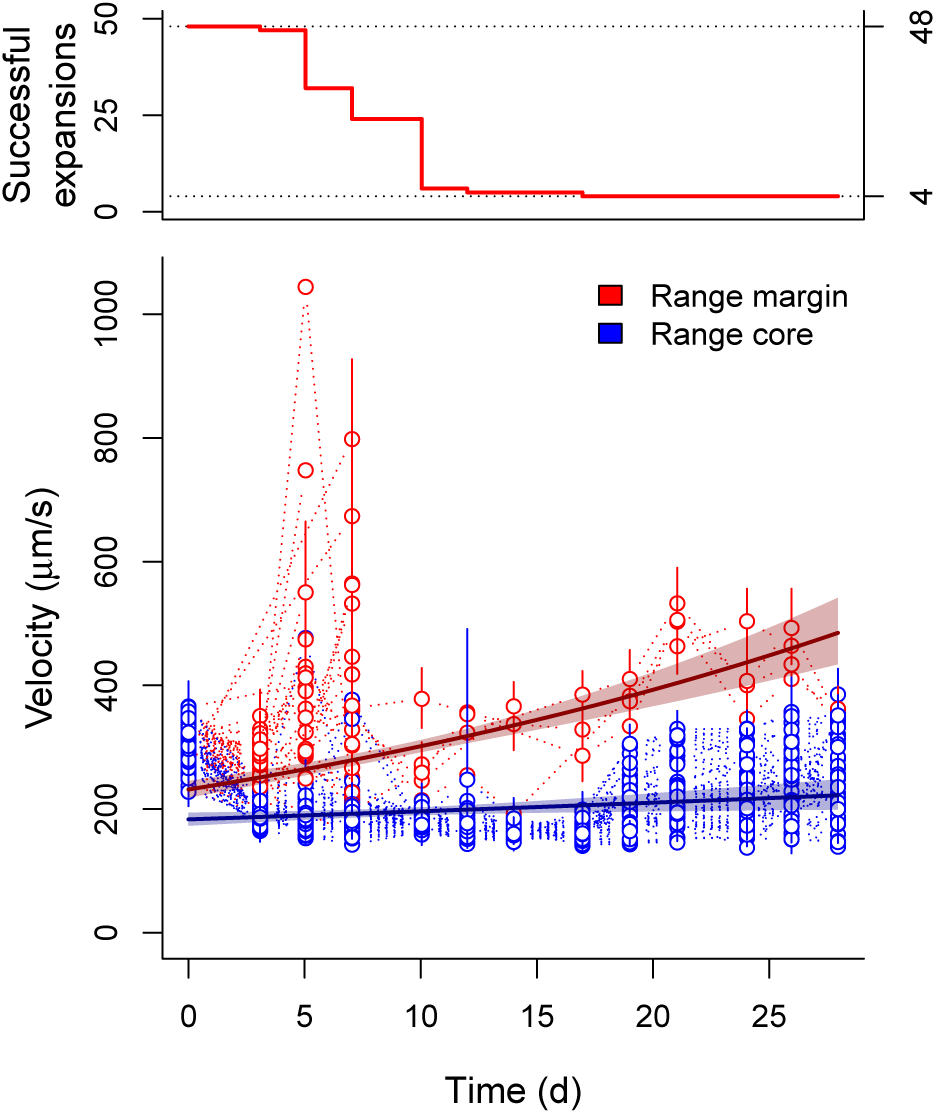
Temporal dynamics of movement velocity during range expansions. While most replicates did not expand their range and went extinct (upper panel), four replicates belonging to three different strains successfully expanded and increased in movement velocity (lower panel). Points and error lines show medians and interquartile ranges of the individual-based velocity data for each replicate separately. We additionally show estimates (solid lines) and confidence intervals (shaded area) obtained from the minimal adequate models correlating velocity and time for range cores (blue) and margins (red).

### Evolution of movement

All four successfully expanding replicates showed increased movement velocity during the experimental evolution phase at the range margin (Fig. 1; LMM; minimal adequate model: time x position; intercept difference core - margin: *t* = −63.65, *d.f.* = 199403, *p* < 0.001; slope margin: *t* = 183.11, *d.f.* = 199403, *p* < 0.001; slope difference core-margin: *t* = −72.38, *d.f.* = 199403, *p* < 0.001). Individuals in populations at the range margin were on average 1.29 times faster than individuals in the range core (comparison of estimates; Fig. 2). Overall, these differences could be shown to be stable both after a three day common garden (approx. 10 to 20 doubling time periods based on the estimates of *r*_0_ in Tab. S1) as well as after an additional month of selection for competitive ability in a common garden setting, thus indicating evolutionary change (Fig. 2; LMM; minimal adequate model: time + position; difference margin - core: *t* = −10.28, *d.f.* = 934, *p* < 0.001; difference end of evolution experiment - common garden 1: *t* = −14.23, *d.f.* = 6, *p* < 0.001; difference common garden 1 - common garden 2: *t* = −14.54, *d.f.* = 6, *p* < 0.001).

**Figure 2:**
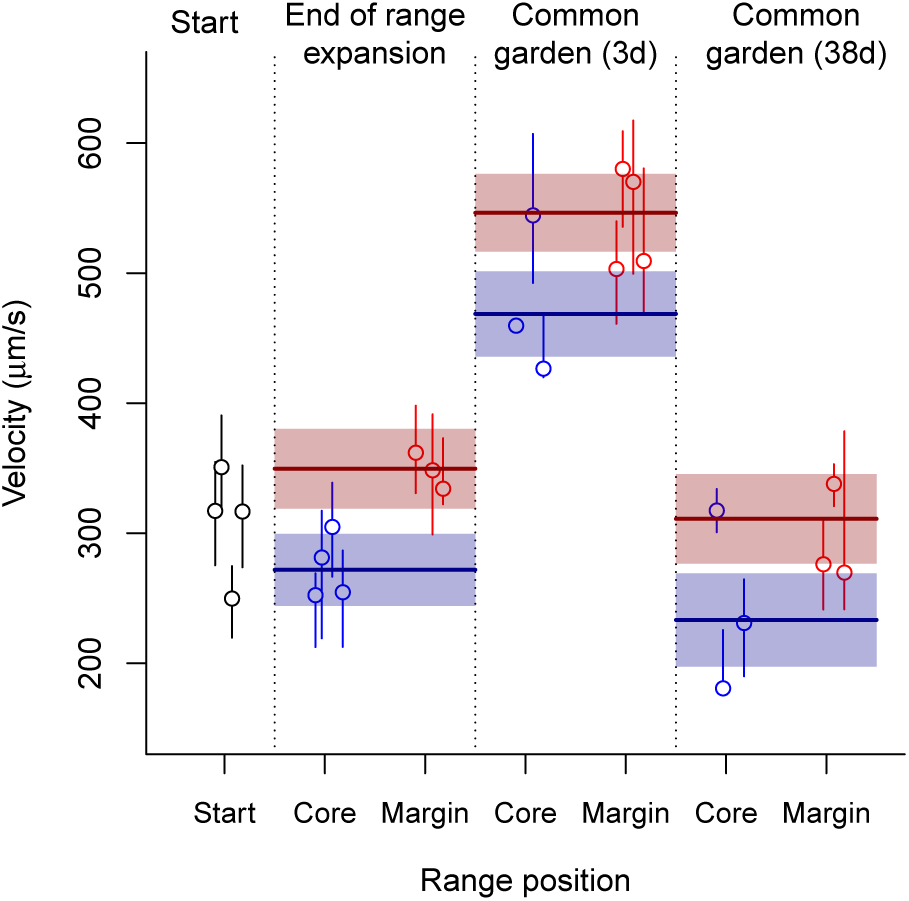
Evolutionary differences in movement velocity between range cores and range margins for the replicates that successfully expanded their range. The increased movement velocity at the range margin at the end of the range expansion (see also Fig. 1) was stable after both the 3-day common garden and the 1-month common garden. Points and error lines show medians and interquartile ranges of the individual-based velocity data for each replicate separately. We additionally show estimates (solid lines) and confidence intervals (shaded area) obtained from the minimal adequate model comparing velocity in rage cores and margins at the three represented time points. Note that velocity at the start of the experiment (black points) is only shown as a visual reference and not included in the statistical analysis.

The successfully expanding replicates did not only show differences in movement after the two common garden periods, which captures evolutionary change only under one specific environmental condition (low densities). In line with our predictions, we also found differences in movement between range core and range margin populations across a large spectrum of population densities (up to two-fold equilibrium density; Fig. 3). The slope of the density-dependent movement reaction norm for range margin populations was negative in comparison to the density-dependent movement reaction norm for range core populations which tended to be slightly positively density-dependent (Fig. 3; LMM on log-transformed data; minimal adequate model: density x history; intercept difference core - margin: *t* = −16.39, *d.f.* = 29608, *p* < 0.001; intercept difference start - margin: *t* = −66.96, *d.f.* = 29608, *p* < 0.001; slope margin: *t* = −4.07, *d.f.* = 29608, *p* < 0.001; slope difference core-margin: *t* = 24.09, *d.f.* = 29608, *p* < 0.001; slope difference start-margin: *t* = 32.67, *d.f.* = 29608, *p* < 0.001).

**Figure 3:**
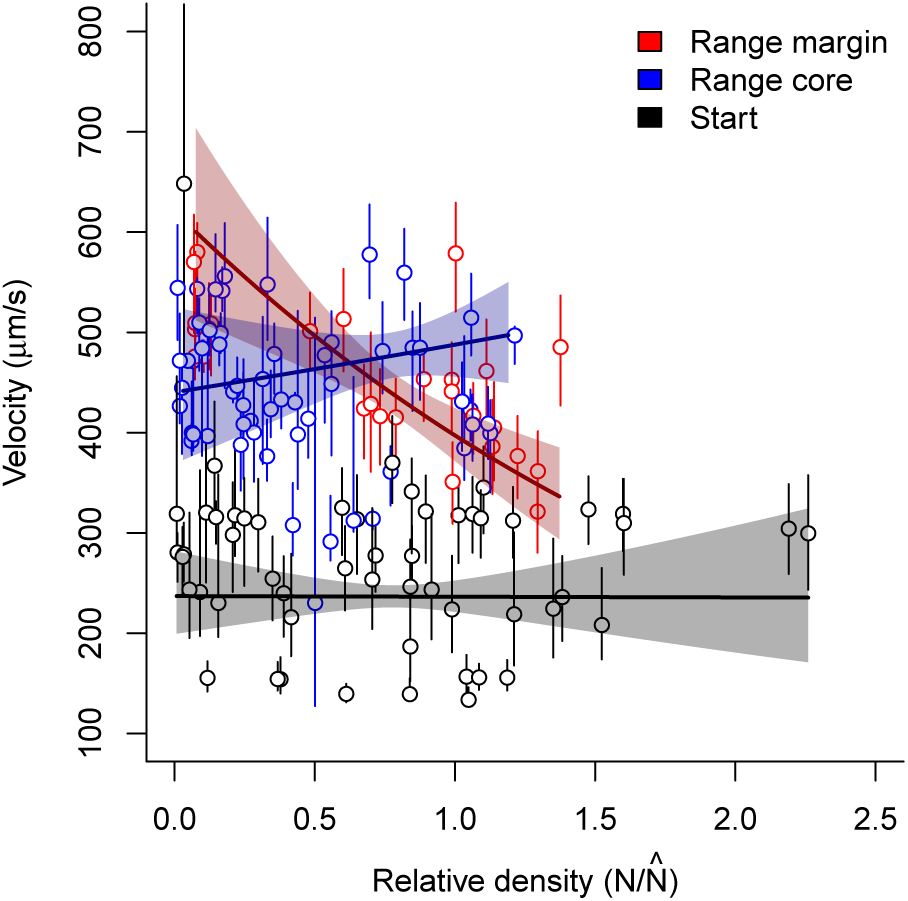
Evolution of density-dependent movement in range cores and at range margins for the replicates that successfully expanded their range. While the shifts in intercept, that is, movement at low densities, recapture the results depicted in Fig. 2, the slope of the density-dependent movement reaction norm is clearly negative at the range margin while it is not significantly different from zero for both range core and starting populations. Points and error lines show medians and interquartile ranges of the individual-based velocity data recorded after the 3-day common garden phase. We additionally show estimates (solid lines) and confidence intervals (shaded area) obtained from the minimal adequate model of the density-dependent movement reaction norm. The x-axis captures the relative density, that is, population size (*N*) divided by the equilibrium population size 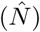 estimated by fitting Eq. 1.

While the differences in intercept observed here recapture the results depicted in Fig. 2, the estimated slopes suggest that density-dependence of movement only evolved at the range margin and did not change in the range core in comparison to the ancestral populations. The estimated slopes for range core and start populations both have confidence intervals that strongly overlap with zero (slope range core: 0.10 (-0.12, 0.32); slope start population: -0.0028 (-0.2166, 0.2113); values represent estimates as well as lower and upper CI for log-transformed data extracted from the minimal adequate model), suggesting no density-dependence. By contrast, the slope of the density-dependent movement reaction norm for the range margin populations is clearly negative (slope range margin: -0.45 (-0.47, -0.42)).

### Evolution of population growth rate and competitive ability

Besides evolutionary changes in density-dependent movement, we could find clear differences in population growth rates after the common garden phase between range core and range margin (Fig. 4). Range margin populations exhibited population growth rates that were on average 3.2 times higher than range core populations (Tukey Post-Hoc test after LMM on log transformed data; core - margin: *z* = −2.82, *N* = 22, *p* = 0.013). Interestingly, while the difference between population growth rates at the range margin and in the starting populations was not significant (Tukey Post-Hoc test after LMM on log transformed data; core - margin: *z* = 0.77, *N* = 22, *p* = 0.72) the difference between range core and start was (Tukey Post-Hoc test after LMM on log transformed data; core - margin: *z* = −2.82, *N* = 22, *p* = 0.013).

**Figure 4:**
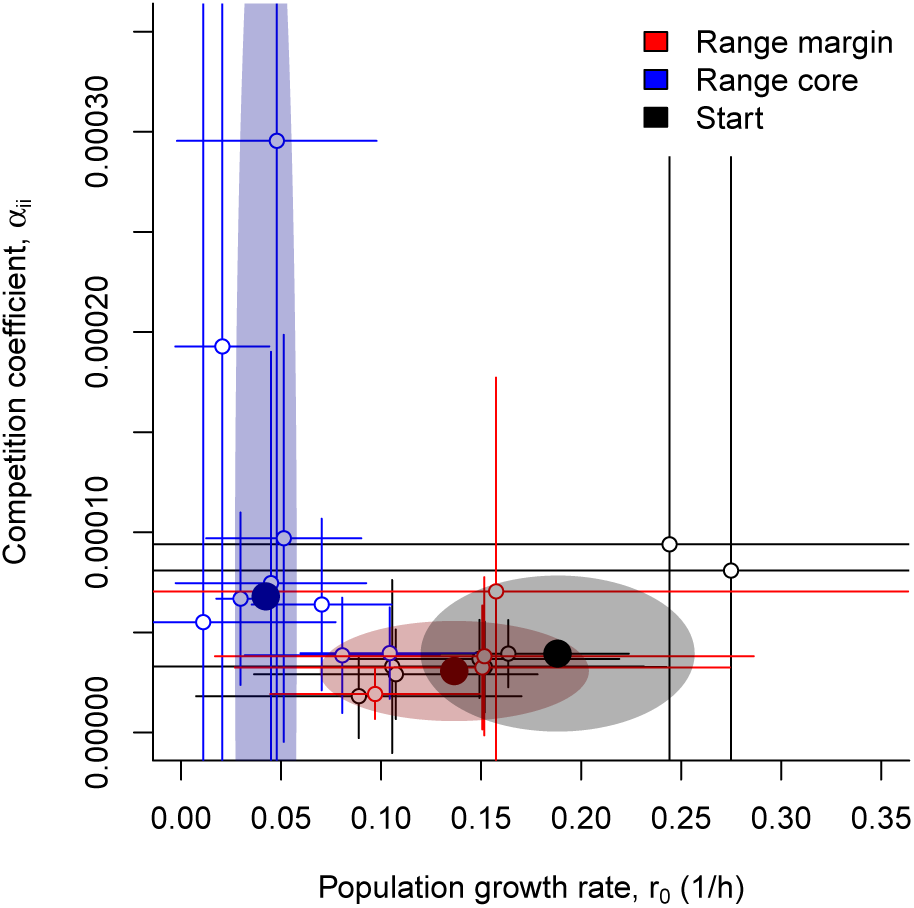
Concurrent evolution of population growth rates (*r*_0_) and competition coefficients (*α*_*ii*_) in range cores and at range margins for the replicates that successfully expanded their range. Range core population showed increased competitive abilities and concurrently decreased population growth rate in comparison to range margins and the starting population. Points and error lines show estimates and confidence intervals of growth rates (*r*_0_) and competition coefficients (*α*_*ii*_) obtained from fitting the logistic growth model (Eq. 1) to data of population growth experiments performed after the 3-day common garden phase. We additionally show estimates (large filled points) and confidence intervals (shaded area) obtained from the minimal adequate model comparing population growth rate and competition coefficient between range cores, margins and the starting populations.

Conversely to population growth rate patterns, range core populations showed a tendency for increased competitive ability in comparison to range margin populations (average increase of 2.2 fold, however with strongly overlapping confidence intervals; Tukey Post-Hoc test after LMM on inverse transformed data; core - margin: *z* = −2.98, *N* = 22, *p* = 0.008). Again, range core populations tended to differ from the starting populations (Tukey Post-Hoc test after LMM on inverse transformed data; core - margin: *z* = 2.28, *N* = 22, *p* = 0.058), while range margin populations did clearly not (Tukey Post-Hoc test after LMM on inverse transformed data; start - margin: *z* = −1.20, *N* = 22, *p* = 0.45).

## Discussion

We investigated evolutionary differentiation between range core and range margin populations using highly controlled and replicated range expansions of both clonal *Tetrahymena thermophila* strains without initial standing genetic variation as well as a mix of all strains. Our experimental results provide evidence of rapidly emerging evolutionary differences in movement reaction norms, population growth rates and competitive ability during range expansions.

As expected, successful range expansions (Fig. 1) occurred relatively rarely (4 out of 48 replicates). All other range margin populations went extinct at some point during the temporally advancing range propagation and ultimately failed to expand their range (Fig. 1). It is interesting to note that we did not observe successful range expansions in the replicates that harboured initial standing genetic variation. Furthermore, we could not identify any common attributes of the strains that successfully expanded their range, which is in good accordance with previous findings highlighting the intrinsically stochastic and variable nature of range expansions, both in an ecological (Melbourne & Hastings, 2009; Giometto *et al.*, 2013) and evolutionary context (Ochocki & Miller, 2017; Weiss-Lehman *et al.*, 2017).

### Range expansions can lead to trait evolution even in the absence of initial standing genetic variation

Evolution of increased dispersal at range margins due to spatial selection (Phillips *et al.*, 2010) and potentially kin competition (Kubisch *et al.*, 2013) has been recognized to impact range expansion dynamics theoretically (Kubisch *et al.*, 2014) as well as in empirical studies (Fronhofer & Altermatt, 2015; Williams *et al.*, 2016; Wagner *et al.*, 2017; Ochocki & Miller, 2017; Weiss-Lehman *et al.*, 2017; Fronhofer *et al.*, 2017). We here explored the relevance of spatial selection for systems that do not initially harbour high amounts of standing genetic variation. Our results may therefore be relevant in the context of invasions of non-native organisms, which, in contrast to expansions of a species’ ranges starting from a large and potentially diverse core habitat, may often start with a colonization event involving one or very few individuals and therefore may exhibit very little standing genetic variation (e.g., Tsutsui *et al.* 2000; but see also Roman & Darling 2007 and Dlugosch & Parker 2008). Given the ecological and economical relevance of invasive species (Pimentel *et al.*, 2000), a better understanding of the underlying evolutionary dynamics is highly valuable.

We show that differentiation between range cores and range margins in movement and other life-history traits (population growth rate and competitive ability) is possible in a relatively short time (13 discrete dispersal and population growth phases; the growth phases correspond to approx. 10 to 20 doubling time periods based on the estimates of *r*_0_ in Tab. S1) even without initial standing genetic variation (Figs. 2 – 4). In our study system rapid evolution in asexually reproducing lines does not seem fundamentally surprising, as mutation rates are relatively high in *T. thermophila*: Brito *et al.* (2010) performed mutation accumulation experiments and report that the observed extinction rate is two to three orders of magnitude higher than in comparable experiments with yeast and bacteria. These high rates are likely due to gain and loss of chromosomes that typically occur during the amitotic division of the macronucleus (Brito *et al.*, 2010).

More generally, our findings fit well into the current invasion literature: For instance, Dlugosch & Parker (2008) discuss the invasive plant *Hypericum canariense* which shows evidence for the evolution of increased growth rates (survival and reproduction) and local adaptation in flowering time in invasive populations despite bottleneck events. Furthermore, genetic diversity does not seem to consistently predict invasion success in nature (Roman & Darling, 2007; Dlugosch & Parker, 2008), which is also consistent with our findings, especially with the failure to expand in the replicates that exhibited standing genetic variation. More generally, that only 4 out of a total of 48 replicates managed to successfully expand their range, fits well with the idea that only a small fraction of introduced and established species, likely orders of magnitude less (the ‘tens rule’; Williamson & Fitter, 1996), become invasive. As Roman & Darling (2007) note, those that do become invasive may not exhibit a general loss of diversity, which is well captured by our results, as those replicates that did expand their range overall showed diversification in life-history traits between core and margin (Fig. 2 and 4).

Besides being applicable to biological invasions, our results and conclusions are potentially relevant to climate- or generally environmental change-driven range expansions. Our experimental procedure basically simulates a shifting window of suitable habitat for the range margin as is often done in theoretical studies (e.g. Henry *et al.*, 2013). Figure 1 highlights that, despite most often leading to extinction, evolutionary change (Fig. 2) can potentially prevent these marginal populations from going extinct (evolutionary rescue; Bell & Gonzalez, 2011; Low-Décarie *et al.*, 2015). Such an interpretation is of course speculative, as we cannot determine whether evolutionary change happened because of the range expansions (spatial selection) or if evolutionary change allowed for the range expansion to happen in the first place.

### Evolution during range expansions changes the density-dependent movement reaction norm

Our work shows that movement and dispersal evolution at expanding range margins does not only impact average movement and dispersal probabilities, but changes the shape of the density-dependent movement reaction norm (Fig. 3). Theory predicts that dispersal should increase even at low densities at expanding range margins, that is, become less positively density-dependent (Travis *et al.*, 2009) because conspecific density looses its value as a cue for context-dependent dispersal. Since potential target patches for range margin populations tend to be empty, dispersal is advantageous already at low densities and an unconditional dispersal strategy will be less costly (Bonte *et al.*, 2012) as energy and time do not have to be invested in information collection and processing.

We find that the starting populations showed weak or no density-dependent movement (Fig. 3). Subsequently, individuals in the range core conserved this density-independent movement strategy, while individuals at the range margin clearly evolved increased movement at low densities (Fig. 3) thereby changing the slope from lightly positive or neutral to negative. Our experimental results therefore substantiate the theoretical predictions of Travis *et al.* (2009) and are in accordance with recent empirical work that suggested a weakening of the slope of density-dependent dispersal function in an insect system (Weiss-Lehman *et al.*, 2017). We can only speculate what led to the overall increase in movement (intercepts) when comparing starting populations and populations after the first common garden (Fig. 2 and 3). Due to the high rates of resource replenishment we chose to avoid evolution in the resources, this shift, which is reversed in the second common garden (Fig. 2), may be due to increased resource availability during the experimental evolution phase.

### Range expansions lead to the evolution of increased competitive ability and decreased population growth rates in range cores

While dispersal is a central and potentially independent life-history trait of major importance for spatial dynamics (Bonte & Dahirel, 2017), other life-history traits, such as reproduction may evolve concurrently or even trade-off with dispersal (e.g. Burton *et al.*, 2010; Fronhofer *et al.*, 2011; Stevens *et al.*, 2014; Fronhofer & Altermatt, 2015). Our results show that not only movement capacities but also reproductive rates and competitive abilities can evolve during range expansions and biological invasions (Fig. 4). Substantiating theoretical predictions (e.g. Burton *et al.*, 2010), we experimentally find higher population growth rates at the range margin in comparison to range core populations. While the slope of the movement reaction norm showed evolutionary change in comparison to the starting population only at the range margin, it is populations in the range core that exhibited lower population growth rates in comparison to populations at the range margin and the starting population and not vice versa. At the same time, we observe higher competitive abilities in range core populations, which hints at a trade-off between competitive ability and population growth rates.

Our results regarding the evolution of population growth and competitive ability, as well as previous findings in a similar system (Fronhofer *et al.*, 2017), are opposed to some recent experimental investigations reporting lower reproduction, respectively investment in population growth at the range margin (Fronhofer & Altermatt, 2015; Weiss-Lehman *et al.*, 2017) or no effect on reproduction (Ochocki & Miller, 2017). While this heterogeneity in results may be linked to differences in resource dynamics in the experiments (Fronhofer & Altermatt, 2015; Fronhofer *et al.*, 2017), it also hints at the potential evolutionary independence of dispersal from other life-history traits (Bonte & Dahirel, 2017). Clearly, more conceptual and experimental research is needed to provide synthesis on these questions.

### Conclusions

We present experimental evidence showing that range expansions lead to the evolution of increased movement at low densities at range margins in comparison to range cores due to evolutionary changes in the density-dependent movement reaction norm. Movement evolved to be negatively density-dependent at the range margin. Additionally, we observed concurrent changes in life-history traits towards lower population growth rates and higher competitive abilities in the range core. Importantly, these evolutionary changes emerged de novo in our experiments and did not require initial standing genetic variation. Our results suggest that evolution of increased movement at range margins may allow individuals to track their climatic niche by adapted dispersal and growth strategies. Furthermore, although rare, evolution can favour the spread of invasive species, even if standing genetic variation is initially low.

## Author contributions

E.A.F. and F.A. designed the research; E.A.F. and S.G. performed the experiments; E.A.F. analysed the data and outlined the manuscript; all authors contributed to writing.

## Acknowledgements

E.A.F. and F.A. thank Eawag and the Swiss National Science Foundation (grant no. PP00P3 150698 to FA) for funding. We are grateful to Felix Moerman for comments on a previous version of the manuscript.

## Supporting Information

E.A. Fronhofer, S. Gut and F. Altermatt:

**Evolution of density-dependent movement during replicated experimental range expansions**

**Figure S1:**
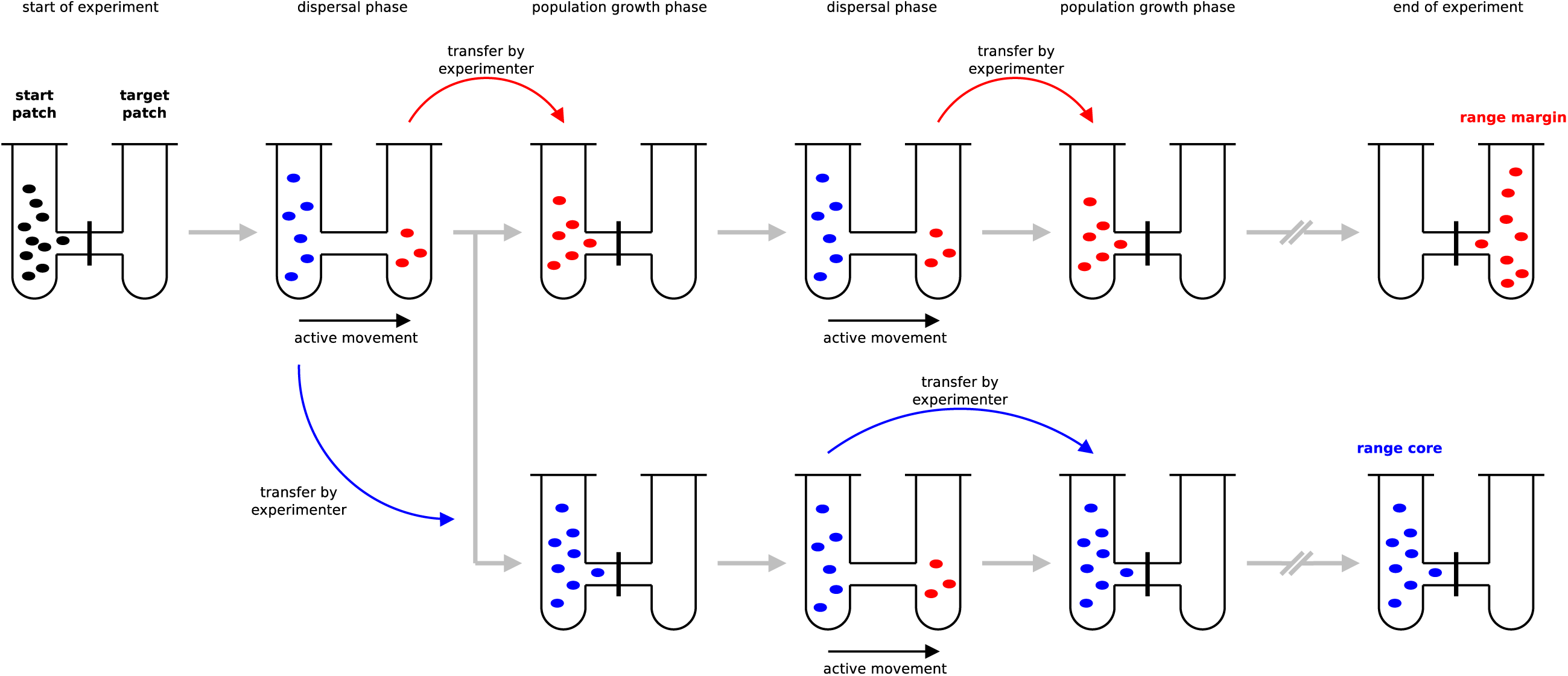
Experimental setup as used here and in Fronhofer & Altermatt (2015). As described in the main text, we effectively tracked range cores (represented in blue) and range margins (represented in red) in a constantly moving time window. While we have chosen this approach for convenience and feasibility and the experimental procedure obviously prevents feedbacks between core and margin populations, our results are not altered if we indeed use long interconnected linear landscapes (Fronhofer *et al.*, 2017). Note that the colors (black for the starting population, blue for the range core and red for the range margin) do not represent any information on genetic background or trait differences. The colors should simply be seen as labels for the spatial location of the populations and are the same as used in the main text.

**Figure S2:**
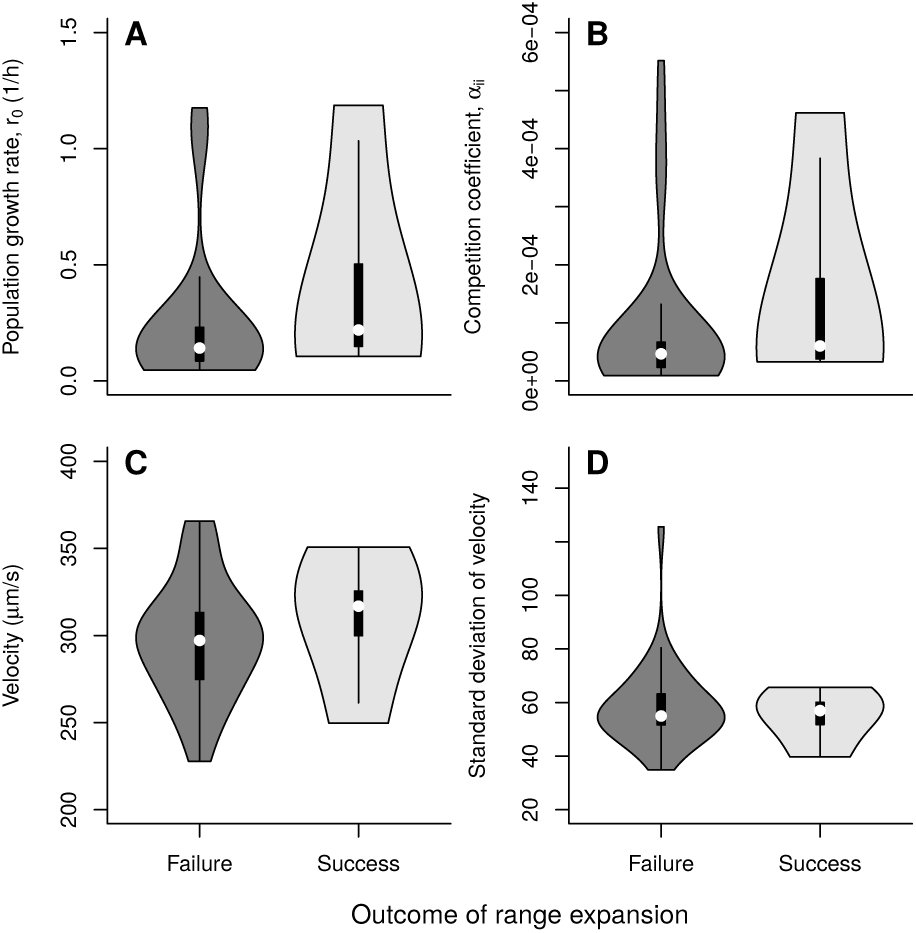
Differences between successful and failed range expansions. The statistical analysis (*r*_0_: LMM on inverse transformed data; *t* = −1.073305, *d.f.* = 25, *p* = 0.2934; *α*_*ii*_: LMM on inverse transformed data; *t* = −1.052423, *d.f.* = 25, *p* = 0.3027; median start velocity: LMM on inverse transformed data; *t* = −0.7424, *d.f.* = 25, *p* = 0.4648; s.d. start velocity: LMM on inverse transformed data; *t* = 1.608692, *d.f.* = 25, *p* = 0.1202) did not detect any differences in strain properties at the start of the experiment between failed an successful range expansions. The violin plots show medians (white dot), the 25th and 75th percentile (box) and the kernel density.

**Table S1:** Strain characteristics: population dynamics. We report median and quartiles of 3 replicates.

| Strain ID | Mating type | Isolate | Population growth rate<br>$r_0$ ($h^{-1}$) | Competition coefficient<br>$\alpha_{ii}$ ($1E-05$ $h^{-1}$) | Equilibrium density<br>$\hat{N}$ ($mL^{-1}$) |
| --- | --- | --- | --- | --- | --- |
| A | I | A | 0.26 (0.17, 0.27) | 6.47 (4.34, 8.38) | 3958 (3327, 4046) |
| B | I | B | 0.16 (0.12, 0.18) | 5.71 (4.16, 6.96) | 2881 (2580, 2895) |
| C | II | A | 0.15 (0.15, 0.21) | 3.68 (3.48, 5.88) | 4055 (3727, 4346) |
| D | II | B | 0.22 (0.18, 0.23) | 5.38 (4.41, 5.90) | 3971 (3902, 3989) |
| E | III | A | 0.06 (0.05, 0.07) | 1.99 (1.88, 2.67) | 2625 (2306, 2850) |
| F | III | B | 0.16 (0.14, 0.62) | 7.27 (6.30, 26.47) | 2180 (2138, 2272) |
| G | IV | A | 0.21 (0.16, 0.64) | 5.37 (5.29, 20.09) | 3079 (2626, 3533) |
| H | IV | B | 0.24 (0.17, 0.72) | 9.40 (6.35, 27.78) | 2598 (2584, 2900) |
| I | V | A | 0.15 (0.11, 0.61) | 5.76 (3.82, 20.84) | 2962 (2754, 3219) |
| J | V | B | 0.21 (0.13, 0.69) | 9.60 (6.19, 32.38) | 2131 (1993, 2167) |
| K | VI | A | 0.11 (0.10, 0.14) | 2.91 (2.36, 3.42) | 4152 (3921, 4525) |
| L | VI | B | 0.06 (0.06, 0.07) | 0.97 (0.95, 1.07) | 6259 (6126, 6425) |
| M | VII | A | 0.13 (0.12, 0.20) | 2.67 (2.52, 4.05) | 4743 (4642, 4975) |
| N | VII | B | 0.13 (0.10, 0.18) | 4.10 (2.75, 5.29) | 3480 (3381, 4141) |

**Table S2:**
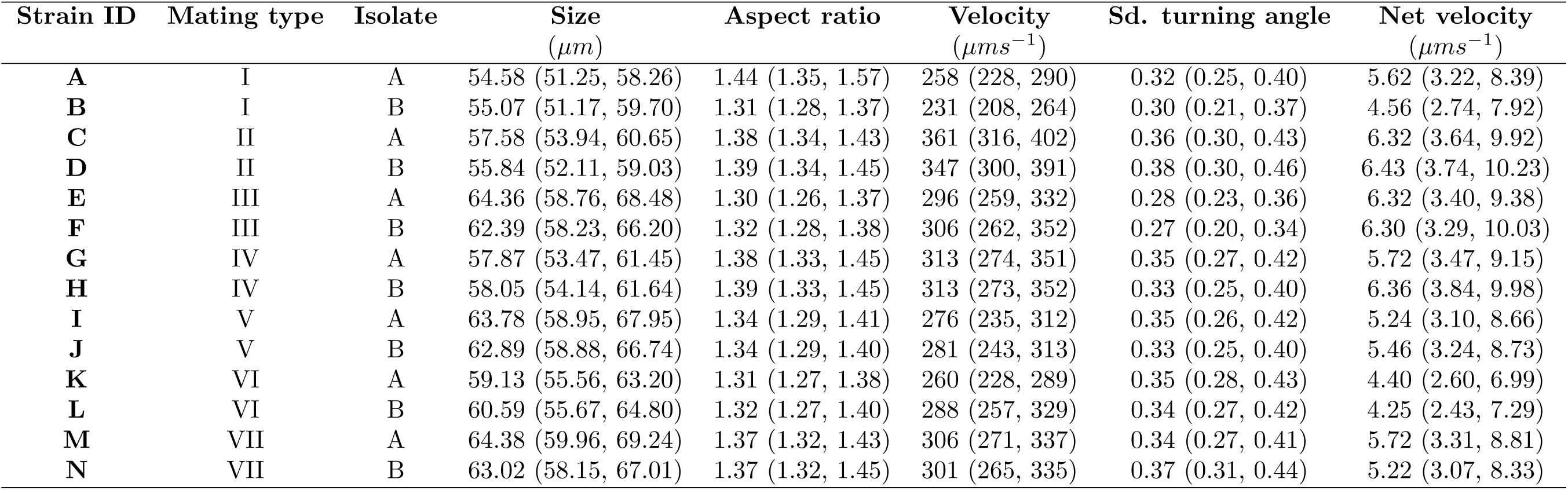
Strain characteristics: morphology and Movement. We report median and quartiles of 3 replicates.

